# Stage-Specific Long Non-coding RNAs in *Cryptosporidium parvum* as Revealed by Stranded RNA-Seq

**DOI:** 10.1101/2020.09.19.304907

**Authors:** Yiran Li, Rodrigo P. Baptista, Adam Sateriale, Boris Striepen, Jessica C. Kissinger

## Abstract

*Cryptosporidium* is a protist parasite that has been identified as the second leading cause of moderate to severe diarrhea in children younger than two and a significant cause of mortality worldwide. *Cryptosporidium* has a complex, obligate, intracellular but extra cytoplasmic lifecycle in a single host. How genes are regulated in this parasite remains largely unknown. Long non-coding RNAs (lncRNAs) play critical regulatory roles, including gene expression across a broad range of organisms. *Cryptosporidium* lncRNAs have been reported to enter the host cell nucleus and affect the host response. However, no systematic study of lncRNAs in *Cryptosporidium* has been conducted to identify additional lncRNAs. In this study, we analyzed a *C. parvum in vitro* strand-specific RNA-seq developmental time series covering both asexual and sexual stages to identify lncRNAs associated with parasite development. In total, we identified 396 novel lncRNAs 86% of which are differentially expressed. Nearly 10% of annotated mRNAs have an antisense lncRNA. lncRNAs also appear to occur most often at the 3’ end of their corresponding sense mRNA. Putative lncRNA regulatory regions were identified and many appear to encode bidirectional promoters. A positive correlation trend between lncRNA and the upstream mRNA expression was observed. Evolutionary conservation and expression of lncRNA candidates was observed between *C. parvum, C. hominis* and *C. baileyi*. Ten *C. parvum* protein-encoding genes with antisense transcripts have *P. falciparum* orthologs that also have antisense transcripts. Three *C. parvum* lncRNAs with exceptional properties (e.g., intron splicing) were experimentally validated using RT-PCR and RT-qPCR. We provide an initial characterization of the *C. parvum* non-coding transcriptome to facilitate further investigations into the roles of lncRNAs in parasite development and host-pathogen interactions.

## Introduction

*Cryptosporidium* is an obligate protist parasite that causes a diarrheal disease called cryptosporidiosis which spreads via an oral-fecal route. Human cryptosporidiosis, mainly caused by *Cryptosporidium parvum* and *Cryptosporidium hominis*, is typically self-limited and causes 1~2 weeks of intense watery diarrhea in people with healthy immune systems. However, the illness may be lethal among the immunocompromised including individuals with AIDS, cancer, and those receiving transplant anti-rejection medications. In recent years, several *Cryptosporidium* species, predominantly *C. hominis*, have been identified as the second most prevalent diarrheal pathogen of infants globally after rotavirus (Bouzid, Hunter et al. 2013, Kotloff, Nataro et al. 2013, Painter, Hlavsa et al. 2015, Platts-Mills, Babji et al. 2015, Sow, Muhsen et al. 2016) and a leading cause of waterborne disease among humans in the United States (Prevention). Thus far, Nitazoxanide, the only FDA-approved drug is not effective for use in infants or those with HIV-related immunosuppression (Amadi, Mwiya et al. 2009) i.e. the most susceptible populations, and no vaccine is available (Amadi, Mwiya et al. 2002).

*Cryptosporidium* has a complex lifecycle in a single host. The *Cryptosporidium* oocyst which is shed in feces is a major extracellular lifecycle stage. It can stay dormant and survive in the environment for months (Drummond, Boano et al. 2018). After ingestion of oocysts through contaminated water or food, sporozoites are released which are capable of invading intestinal epithelial cells where both asexual and sexual replication occur. Following invasion, sporozoites develop into trophozoites and undergo asexual replication to generate type I meronts and type II meronts. Type I meronts are thought to be capable of reinvading adjacent cells generating an asexual cycle (Fayer 2008), while Type II meronts contribute to the formation of microgamonts (male form) or macrogamonts (female form) to complete the sexual stages (Bouzid, Hunter et al. 2013). Conventional monolayer cell culture does not permit completion of the life cycle much beyond 48 hours post-infection (hpi) (**Figure 1A**), for as of yet poorly understood reasons but gametogenesis does occur (Tandel, English et al. 2019). The lack of *in vitro* culture has historically impeded the development of new drugs and vaccines for this medically important parasite. Recently, there have been several breakthroughs including genetic manipulation (Vinayak, Pawlowic et al. 2015, Sateriale, Pawlowic et al. 2020) and lifecycle completion. Several promising approaches have been developed including using a cancer cell line as host (Miller, Josse et al. 2018), biphasic and three-dimensional (organoid) culture systems, (Morada, Lee et al. 2016, DeCicco RePass, Chen et al. 2017, Heo, Dutta et al. 2018, Cardenas, Bhalchandra et al. 2020), hollow fiber technology (Yarlett, Morada et al. 2020), and air-liquid interface (ALI) cultivation system (Wilke, Funkhouser-Jones et al. 2019, Wilke, Wang et al. 2020). These breakthroughs are enabling better, much needed, studies of the parasite’s full life cycle. A better understanding the conditions and regulatory processes necessary for *Cryptosporidium* development are essential and will prove beneficial for the identification of drug and vaccine targets.

**Figure 1.**
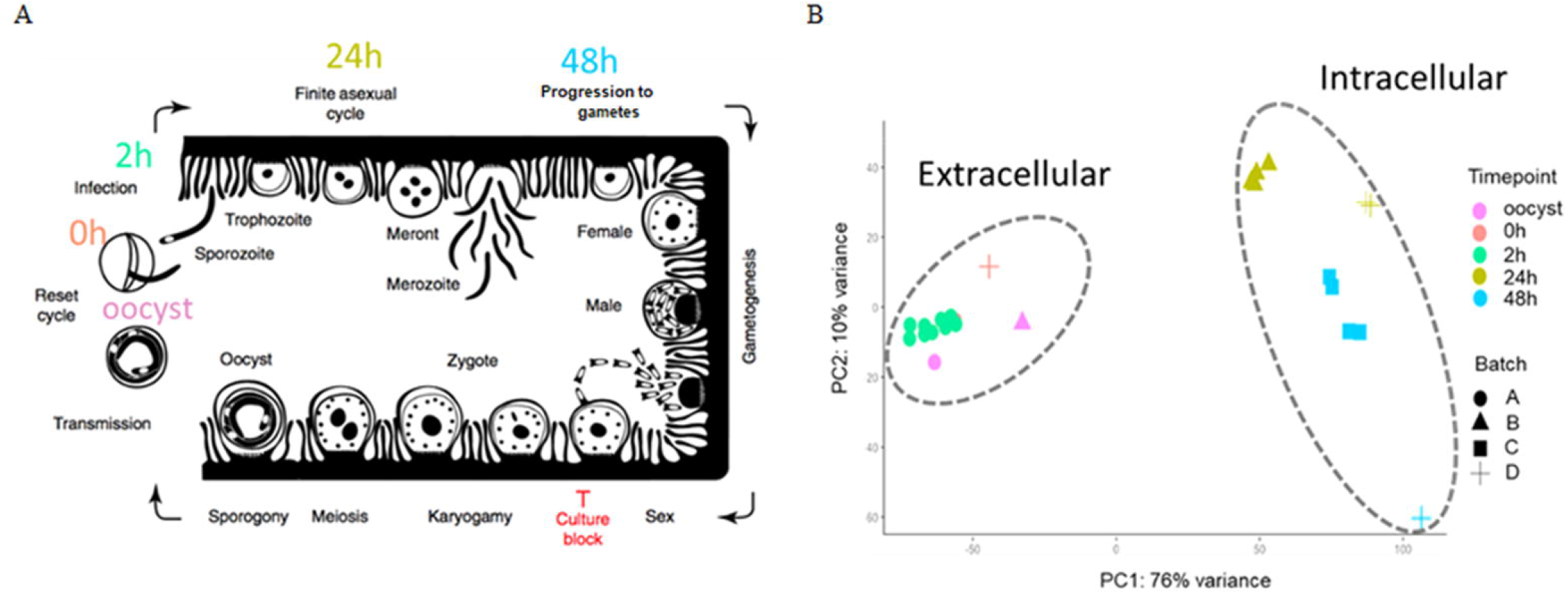
The 33 RNA-Seq datasets used for expression analysis. A) The time points indicate when RNA-Seq samples were collected and the associated *C. parvum* life cycle stage. The schematic model of the *C. parvum* life cycle is reproduced from (Tandel, English et al. 2019). B) Principal component analysis of 33 *C. parvum* transcriptomes. The analysis is based on the normalized expression level (VST) of the *C. parvum* mRNA and predicted lncRNA genes. Samples collected from different time points are indicated by colors. Various projects/batches are represented by shapes. Batch A includes sample IDs 3-9 without host cells, Batch B includes sample IDs 1, 23-26 with host cell type of IPEC, Batch C includes sample IDs 27-30 26 with host cell type of MDBK, Batch D contains sample IDs 2, 20-22, 31-33 with host cell type of HCT-8 (Table 1).

The first genome sequence of *C. parvum* was published in 2004 with a genome size of ~9 Mb and ~3800 protein-encoding genes annotated (Abrahamsen, Templeton et al. 2004). Since this milestone, our understanding of the parasite and its biology have progressed remarkably. Early *in vitro* transcriptome analyses using semi-quantitative RT-PCR over a 72 h post-infection (pi) time course during *in vitro* development revealed complex and dynamic gene expression profiles. Adjacent genes are not generally co-regulated, despite the highly compact genome (Mauzy, Enomoto et al. 2012). Under UV irradiation, *C. parvum* oocysts have shown a vital stress-induced gene expression response according to microarray data (Zhang, Guo et al. 2012). mRNA expression related to gametocyte and oocyst formation were studied using RNA sequencing of sorted cells (Tandel, English et al. 2019). Yet, little is known about the regulation of key developmental transitions. How the parasite regulates gene expression in order to invade hosts, amplify, evade the immune system and interact with their host awaits further discovery.

Most canonical eukaryotic enhancer proteins are not detected in Apicomplexa, the phylum that *Cryptosporidium* belongs to (Iyer, Anantharaman et al. 2008), except for the transcriptional activators Myb and zinc finger proteins C2H2 (Cys2His2) and two additional transcription factor families. Instead, an expanded family of apatela-related transcription factors, the AP2 family of proteins (ApiAP2), appear to be the predominant transcription factors in this phylum, including *Cryptosporidium* (Oberstaller, Pumpalova et al. 2014, Jeninga, Quinn et al. 2019). AP2 domains in *C. parvum* are reported to have reduced binding diversity relative to the malaria parasite *Plasmodium falciparum* and proposed to possess less dominancy in transcriptional regulation in *C. parvum* (Campbell, De Silva et al. 2010, Yarlett, Morada et al. 2020). It has been proposed that *C. parvum* is less reliant on ApiAP2 regulators in part because it utilizes E2F/DP1 transcription factors, which are present in *Cryptosporidium* while absent in other studied apicomplexans (Templeton, Iyer et al. 2004, Yarlett, Morada et al. 2020). Based on the similarity of gene expression profiles, it has been suggested that the number of co-expressed gene clusters in *C. parvum* is somewhere between 150 and 200, and putative ApiAP2 and E2F/DP1 *cis*-regulatory elements were successfully detected in the upstream region of many co-expressed gene clusters (Oberstaller, Joseph et al. 2013). Additionally, low levels of DNA methylation in oocysts has been reported in several *Cryptosporidium spp*. (Gong, Yin et al. 2017), suggesting the requirement of additional regulatory mechanisms. At the level of post-transcriptional regulation, the RNA interference (RNAi) pathway, which plays a crucial role in gene silencing in most eukaryotes, is considered to be missing in *Cryptosporidium* due to the lack of identifiable genes encoding the microRNA processing machinery or RNA-induced silencing complex (RISC) components (Keeling 2004). There much that remains to be discovered with respect to regulation of gene expression in *Cryptosporidium*.

Long non-coding RNAs (lncRNAs) are transcripts without significant protein-encoding capacity that are longer than 200 nt. In eukaryotes, lncRNAs play critical regulatory roles in gene regulation at multiple levels, including transcriptional, post-transcriptional, chromatin modification and nuclear architecture alterations (Marchese, Raimondi et al. 2017). In humans, 3,300 long intergenic ncRNAs (lincRNAs) were analyzed using chromatin state maps, and ~20% of these RNAs are bound to the polycomb repressive complex (PCR2, a complex with histone methyltransferase activity) (Khalil, Guttman et al. 2009). Most lncRNAs share many characteristics of mRNAs, such as RNA polymerase II-mediated transcription, a 5’ 7-methylguanosine cap and a 3’ poly(A) tail [6]. The expression of lncRNAs is usually more tissue- or time-specific than mRNA expression (Necsulea, Soumillon et al. 2014, Tsoi, Iyer et al. 2015). lncRNA sequences are not well conserved across species, but their structure could be conserved due to functional constraints (Ulitsky, Shkumatava et al. 2011, Diederichs 2014). By forming hybrid structural complexes such as RNA-DNA hybrid duplexes or RNA-DNA triplexes, lncRNAs can recruit or scaffold protein complexes to facilitate localization of protein machinery to specific genome target sites (Li, Mo et al. 2016). lncRNAs play important roles in regulating occurrence and progression of many diseases. After infected by *C. baileyi*, significant expression changes have been observed in the host (Ren, Fan et al. 2018). The mis-regulation of lncRNAs in multi-cellular eukaryotes has been shown to cause tumorigenesis (Chakravarty, Sboner et al. 2014), cardiovascular diseases (Tang, Mei et al. 2019), and neurodegenerative dysfunction (Zhang, Luo et al. 2018) and thus can be used as diagnostic biomarkers.

Taking advantage of sequencing technologies, numerous lncRNA candidates have been detected in apicomplexans, some with proven regulatory potential during parasite invasion and proliferation processes. These discoveries have ushered in a new era in parasite transcriptomics research (Liao, Shen et al. 2014, Siegel, Hon et al. 2014, Broadbent, Broadbent et al. 2015, Ramaprasad, Mourier et al. 2015, Filarsky, Fraschka et al. 2018). In *P. falciparum*, lncRNAs are critical regulators of virulence gene expression and associated with chromatin modifications (Vembar, Scherf et al. 2014). Likewise, an antisense lncRNA of the gene *gdv1* was shown to be involved in regulating sexual conversion in *P. falciparum* (Filarsky, Fraschka et al. 2018). In *C. parvum*, putative parasite lncRNAs were found to be delivered into the host nucleus, some of which were experimentally proven to regulate host genes by hijacking the host histone modification system (Ming, Gong et al. 2017, Wang, Gong et al. 2017, Wang, Gong et al. 2017). The importance of lncRNA in *C. parvum* has been demonstrated, but no systematic annotation of lncRNA has been conducted. The systematic identification of lncRNAs will increase the pool of candidate regulatory molecules thus ultimately leading to increased knowledge of the developmental gene regulation in *C. parvum* and control of parasite interactions with its hosts.

In this study, we developed and applied a computational pipeline to systematically identify new lncRNAs in the *C. parvum* genome. We used a set of stranded RNA-seq data collected from multiple lifecycle stages that cover both asexual and sexual developmental stages. We conducted a systematic analysis of lncRNA that includes sequence characteristics, conservation, expression profiles and expression relative to neighboring genes. The results provide new insights into *C. parvum’s* coding potential and suggest several areas for further research.

## Methods and Materials

### RNA-Seq data pre-processing/cleaning

RNA-Seq datasets were downloaded from the NCBI Sequence Read Archive (SRA) and European Nucleotide Archive database (ENA). Detailed information on SRA accession numbers and Bioprojects are listed in **Table 1** and **Supplementary Table S1**. FastQC-v0.11.8 was used to perform quality control of the RNA-Seq reads. Remaining adapters and low-quality bases were trimmed by Trimmomatic-v-0.36 (Bolger, Lohse et al. 2014) with parameters: Adapters:2:30:10 LEADING:20 TRAILING:20 SLIDINGWINDOW:4:25 MINLEN:25. All reads were scanned with a four-base sliding window and cut when the average Phred quality dropped below 25. Bases from the start and end were cut off when the quality score was below 20. The minimum read length was set at 25 bases. The processed reads are referred to as cleaned reads in this work.

**Table 1.** Mapping statistics of RNA-Seq datasets

| ID* | Time point | Host cells | Condition |
| --- | --- | --- | --- |
| *C. parvum* |  |  |  |
| 1B | oocyst | NA | extracellular |
| 2D | oocyst | NA | extracellular |
| 3A | 0h | NA | extracellular |
| 4A | 0h | NA | extracellular |
| 5A | 0h | NA | extracellular |
| 6A | 0h | NA | extracellular |
| 7A | 0h | NA | extracellular |
| 8A | 0h | NA | extracellular |
| 9A | 0h | NA | extracellular |
| 10A | 2h | NA | extracellular |
| 11A | 2h | NA | extracellular |
| 12A | 2h | NA | extracellular |
| 13A | 2h | NA | extracellular |
| 14A | 2h | NA | extracellular |
| 15A | 2h | NA | extracellular |
| 16A | 2h | NA | extracellular |
| 17A | 2h | NA | extracellular |
| 18A | 2h | NA | extracellular |
| 19A | 2h | NA | extracellular |
| 20D | 24h | HCT-8 | intracellular |
| 21D | 24h | HCT-8 | intracellular |
| 22D | 24h | HCT-8 | intracellular |
| 23B | 24h | IPEC | intracellular |
| 24B | 24h | IPEC | intracellular |
| 25B | 24h | IPEC | intracellular |
| 26B | 24h | IPEC | intracellular |
| 27C | 48h | MDBK | intracellular |
| 28C | 48h | MDBK | intracellular |
| 29C | 48h | MDBK | intracellular |
| 30C | 48h | MDBK | intracellular |
| 31D | 48h | HCT-8 | intracellular |
| 32D | 48h | HCT-8 | intracellular |
| 33D | 48h | HCT-8 | intracellular |
| *C. hominis* |  |  |  |
| 34A | oocyst | NA | extracellular |
| 35A | oocyst | NA | extracellular |
| 36F | oocyst | NA | extracellular |
| 37F | oocyst | NA | extracellular |
| *C. baileyi* |  |  |  |
| 38G | Oocyst/sporozoite | NA | extracellular |

\*Batch with various host cell types and parasites from different projects are indicated with the sample IDs. IDs designated with A, C and E: (Matos, McEvoy et al. 2019); B: (Mirhashemi, Noubary et al. 2018); D: (Tandel, English et al. 2019); 36F: SRR1183950; 37F: SRR1183934; 38G: SRR1183952.
Cell line synonyms/origin: HCT-8: Human intestine; IPEC: Intestinal Porcine Epithelial Cell line; MDBK: Bovine kidney

### Read mapping and transcript assembly

Cleaned reads from each sample were mapped to the reference genome sequence for *Cryptosporidium parvum* IOWA-ATCC (Baptista et al. in prep) downloaded from CryptoDB v46 (https://cryptodb.org/cryptodb/) using the mapping tool HISAT2-v2.1.0 (Kim, Langmead et al. 2015) with maximum intron length (--max-intronlen) set at 3000, and remaining parameters as default. Uniquely mapped reads were selected for further study using SAMtools-v1.10 (view -q 10) (Li, Handsaker et al. 2009). StringTie-v2.0.6 (Pertea, Pertea et al. 2015) was used to reconstruct transcripts using the reference annotation guided method (--rf -j 5 -c 10 -g 1). At least five reads were required to define a splice junction. A minimum read coverage of 10 was used for transcript prediction. Only overlapping transcript clusters were merged together. The stranded library types were all “fr-firststrand”. Transcripts with FPKM lower than three were removed. The transcriptome assemblies from multiple samples were merged into one master transcript file using TACO-v0.7.3 with default settings (Niknafs, Pandian et al. 2017).

### lncRNA prediction

Transcripts that overlapped with currently annotated mRNAs in the *C. parvum* IOWA-ATCC annotation in CryptoDB v46 with coverage >70% on the same strand were removed using BEDTools-v2.29.2 (Quinlan and Hall 2010). The remaining transcripts were examined for coding potential using the online tool Coding Potential Calculator (CPC) v0.9 (Kong, Zhang et al. 2007). Transcripts considered as “coding” by CPC were removed. Potential read-through transcripts resulting from transcription of neighboring mRNAs were removed using two criteria: 1) The transcript was <50 bp from the upstream coding region of another gene on the same strand. 2) The transcript was always transcribed together with the upstream mRNA on the same strand. Finally, the remaining transcripts which occurred in >2 RNA-Seq samples were kept as putative lncRNA candidates for further studies.

### Transcriptome data normalization and identification of differentially expressed genes

The raw read counts for both mRNA genes and predicted lncRNAs were calculated using HTSeq-v0.12.4 (Anders, Pyl et al. 2015). All genes were filtered to require > 50 reads in at least three samples. Variance stabilizing transformation (VST) from DESeq2-v1.28.1 (Love, Huber et al. 2014) was used to normalize the expression between samples. Principal components analysis (PCA) of the RNA-Seq samples was performed using the resulting VST values. VST values for each of the mRNA and lncRNA candidates were visualized using the R package pheatmap-v1.0.12 (https://github.com/raivokolde/pheatmap). The K-mean approach in pheatmap was applied to cluster genes based on the expression data.

Differentially expressed genes including mRNAs and lncRNAs were analyzed by EdgeR-v3.30.3 (Robinson, McCarthy et al. 2010). The expression time-points compared were between oocyst/sporozoites (oocysts, time point 0 h and sporozoites, 2 h), asexual stage (time point 24 h) and mixed asexual/sexual stage (selected samples from 48 h time point in which gametocyte marker genes are clearly expressed). The sex marker gene: cgd6_2090 encodes *Cryptosporidium* oocyst wall protein-1 (COWP1) produced in female gametes and cgd8_2220 encodes the homolog of hapless2 (HAP2) a marker of male gamonts (Tandel, English et al. 2019). A generalized linear model (GLM) approach was used for differential expression hypothesis testing. P-values were adjusted by the false discovery rate (FDR). Significant differentially expressed genes were declared at a log2-fold change ≥ 1.5 and an FDR < 0.05.

### Expression correlation

The expression correlation analysis between predicted lncRNAs and mRNAs was conducted with normalized expression data from all 33 samples (**Table 1**) of *C. parvum* using the Pearson test. P-values were adjusted by FDR.

### Upstream motif analysis

MEME v5.0.0 (Bailey and Elkan 1994) was used to discover motifs that may be present upstream of putative lncRNAs. For the promoter motif search, we extracted the 100 bp of (+) strand sequence upstream of the predicted lncRNAs and searched both strands using MEME for motifs with length six to 50 bp. The parameters were: -dna -mod anr -nmotifs 6 -minw 6 -maxw 50 -objfun classic -markov_order 0.

### Evolutionary conservation

RNA-Seq datasets from *C. hominis* and *C. baileyi* oocysts were mapped to reference genome sequences for *C. hominis* 30976 and *C. baileyi* TAMU-09Q1, respectively, downloaded from CryptoDB v46 (https://cryptodb.org/cryptodb/). Mapped reads were assembled into transcripts using the same methods mentioned above. tblastx from NCBI BLAST v 2.10.0 was used to search for conserved antisense lncRNA candidates among *C. parvum*, *C. hominis* and *C. baileyi*, with parameter of E-value:1e-5 and best hit retained. To assess conservation among other apicomplexans, *P. falciparum*, antisense and associated sense mRNA information were retrieved from (Broadbent, Broadbent et al. 2015). *P. falciparum* orthologs of the sense mRNAs in *C. parvum* were retrieved from OrthoMCL DB v6.1 (Li, Stoeckert et al. 2003). If antisense lncRNAs were detected between *P. falciparum* and *C. parvum* orthologs, the lncRNAs between *P. falciparum* and *C. parvum* were considered conserved.

### RT-qPCR validation

We designed PCR primers using the PrimerQuest Tool from IDT (https://www.idtdna.com/pages/tools/primerquest) (**Supplementary Table S2**) to use for expression validation and exon structure confirmation of select lncRNA candidates using RT-PCR and qPCR. RNA was extracted from oocysts and provided by Boris Striepen. The cDNA for each sample was reverse-transcribed using the iScript™ cDNA Synthesis Kit (Bio-Rad, Hercules, CA) from 1 □g of input RNA. The resulting cDNAs were used as templates for PCR amplification and qPCR detection. Strand-specific primers were designed to amplify antisense RNAs. The 18S rRNA gene of *C. parvum* was used as positive control and samples without RNA or primer were used as a negative control. Each RT-PCR reaction contained 1ul cDNA, 2 □1 primer mix (10□M), 2 □1 water and 5□1 MyTaq™ HS Mix (Bioline). RT-PCR was performed in the following conditions: 35 cycles of 15 seconds at 95°C, 30 seconds at 64°C. Then the RT-PCR products were run on a 2% agarose gel. The cDNA was also subjected to qPCR with All-in-One qPCR Mix (QP001; GeneCopoeia, Rockville, MD, USA) using the Mx3005P qPCR system (Agilent Technologies, Santa Clara, CA, USA). All reactions, including no-template controls, were run in triplicate. Following amplification, the CT values were determined using fixed threshold settings. lncRNA expression was normalized to 18S rRNA expression.

### Data availability

The GenBank accession records CP044415-CP044422 have been updated to include annotation of the lncRNAs identified in this study. These data have also been submitted to CryptoDB.org.

## Results

### Mapping statistics of stranded RNA-Seq data

To identify and investigate the expression profile of lncRNAs in *C. parvum* during parasite development, we searched for stranded RNA-Seq data sets from *Cryptosporidium* available in public databases. Due to the small volume of *Cryptosporidium* relative to the host cells, RNA-Seq data usually suffer from high host contamination. In this study, we selected samples that had more than 100k *Cryptosporidium* reads generated from the Illumina platform mapped to the reference genome sequence to reduce bias mostly arising from sequencing platform and sequencing depth. In total, 38 stranded-RNA-seq data sets which originated from 33 *C. parvum* samples, four *C. hominis* samples and one *C. baileyi* sample were selected for further analysis. The details and mapping statistics of each sample are shown in **Table 1** and (**Supplementary Table S1**). The *C. parvum* samples represented five time points: oocyst, 0 h (sporozoites immediately after oocyst excystation), 2 h (2-h incubation in the medium (Matos, McEvoy et al. 2019)), 24 h (24-h post host cell infection) and 48 h (48-h post host cell infection). The 24 h and 48 h samples were derived from different types of host cells (see details in **Table 1** and **Supplementary Table S1**). The *C. hominis* samples and one *C. baileyi* sample were obtained from oocysts.

### Identification and characteristics of lncRNAs

We began assembly of the *C. parvum* transcriptome using mapped RNA-Seq reads of samples in NCBI Bioproject PRJNA530692 with a high sample quality. However, *C. parvum* has an extremely compact genome sequence. As calculated from the *C. parvum* IOWA-ATCC annotation, the average intergenic distance between the stop and start codon boundaries of neighboring genes is only 504 bp and this distance must also include promoter and UTR regions. The average length of an annotated *C. parvum* ATCC mRNA coding sequence, CDS, is 1802bp. The high gene density and the short distance between genes make it difficult to set UTR boundaries using short-read sequencing data as transcripts overlap and become merged. Transcriptome assembly using RNA-Seq without genome and reference annotation guidance would lead to a high chimerism rate. Thus, we used reference annotation to guide the assembly process and set parameters to minimize the number of artificially fused transcripts. We then used the program TACO v0.7.3 (Multi-sample transcriptome assembly), which employs change point detection to break apart complex loci to lower the number of fused transcripts to obtain a non-redundant master transcriptome from all samples, resulting in 5818 transcripts in total. Of these, 4846 transcripts overlapped with an mRNA on the same strand and thus, were removed. Transcripts which were <200 bp or only detected in a single sample were removed to improve the lncRNA prediction quality. Transcripts that were considered as “coding” by the Coding potential analysis tool CPC were filtered out. To identify and remove potential read-through transcripts, predicted transcripts that were located closer than 50 bp from the coding region of the upstream gene on the same strand and always transcribed together with the upstream mRNA were removed (**Figure 2A**).

**Figure 2.**
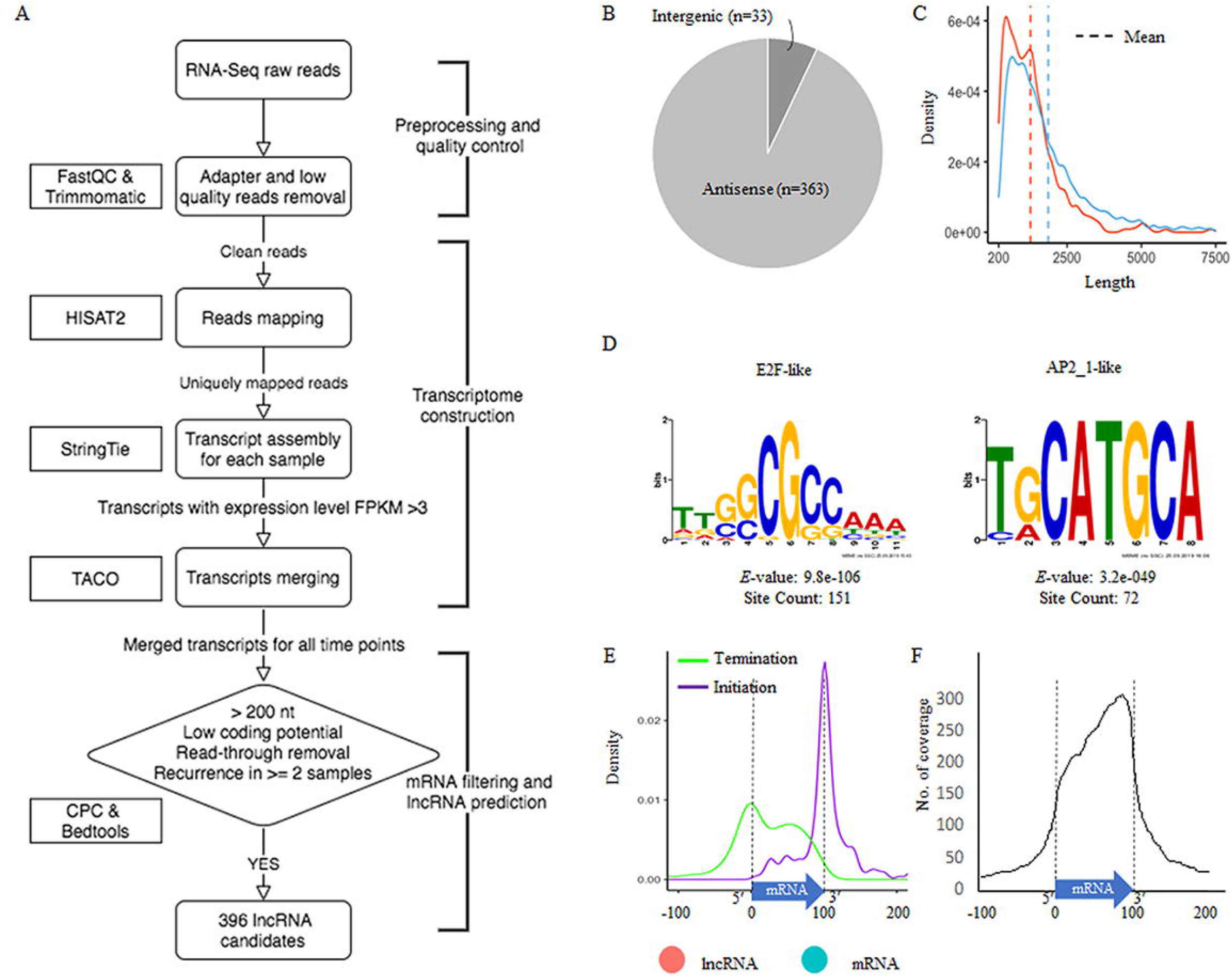
Prediction and characterization of lncRNAs in *C. parvum*. A) Pipeline of lncRNA prediction. B) Genomic location of predicted lncRNAs. C) The distribution of transcript length of mRNA genes and lncRNA candidates. D) Enriched upstream motifs within 100 bp, the same strand (+) of lncRNA candidates E) Antisense transcription initiation and termination position relative to the sense gene body (normalized to 0-100). F) Abundance and position of sense gene body (normalized to 0-100) covered by antisense transcription

In total, 396 transcripts, primarily located antisense to an mRNA (**Figure 2B**), were selected as lncRNA candidates for further analysis. Most of the lncRNAs we detected consist of a single exon however five lncRNAs contain introns. This is consistent with the low-intron rate in *Cryptosporidium*. Additional introns are expected to emerge with deeper RNA-Seq data since many lncRNAs have low expression levels. The average length of the lncRNAs and mRNAs is transcripts 1267 bp and 1866 bp (including UTRs), respectively (**Figure 2C**). When compared to mRNAs, one of the most distinguishing features of lncRNAs is the low Open Reading Frame (ORF) coding potential relative to the transcript length. To understand their biogenesis, we searched for potential promoter motifs within 100 bp of the (+) strand upstream from all 396 lncRNAs. This analysis returned five significant motifs (E-value<0.001), the top two, which were also the most dominant motifs, are related to known *Cryptosporidium* transcriptional factor binding motifs for mRNA genes. It included the E2F/DP1 (5’-[C/G]GCGC[G/C]-3’) and ApiAP2_1 (5’-BGCATGCAH-3’) motifs (**Figure 2D**). This suggests that lncRNA transcriptions have the potential for being regulated independently during parasite development.

We further investigated the relative location of the antisense transcripts relative to the mRNA gene body, and we found that many of lncRNAs’ initiation and termination sites are located close to the gene body boundaries, especially the start site of the antisense lncRNA transcript (assembled transcript including untranslated regions, UTRs) (**Figure 2E**). This trend is most apparent with the lncRNA initiation site. When looking at the coverage of the antisense transcript on mRNA, the *C. parvum* lncRNA antisense expression has a bias towards the 3’ end of the mRNA transcript (**Figure 2F**). This property has also been reported in other organisms, including the malaria pathogen *Plasmodium falciparum* (Lopez-Barragan, Lemieux et al. 2011, Siegel, Hon et al. 2014, Broadbent, Broadbent et al. 2015).

### The *Cryptosporidium* transcriptome varies developmentally and by host

Before profiling the expression of lncRNA candidates, we first compared the transcriptomes of the 33 *C. parvum* RNA-Seq samples by principal component analysis (PCA) based on the normalized mRNA and lncRNA gene expression level of each sample (**Figure 1B**). Extracellular stages, including oocyst and sporozoites from 0 h and 2 h, are differentiated from intracellular stages, including 24 h and 48 h. The transcriptomes of intracellular stages were demonstrated to be more heterogeneous, while extracellular samples formed a relatively compact cluster. This observation was consistent with a previous transcriptome study of *C. parvum* oocysts and intracellular stages (Matos, McEvoy et al. 2019). At time points 24 h and 48 h, different host cells and laboratory procedures were used, which could contribute to the distance observed between samples from the same time point.

To further explore whether sexual commitment was initiated in all 48 h samples, we profiled the transcriptome of marker genes cgd6_2090 and cgd8_2220 (**Supplementary Figure 1**). cgd6_2090 encodes the *Cryptosporidium* oocyst wall protein-1 (COWP1) which is produced in female gametes (Tandel, English et al. 2019); cgd8_2220 encodes the homolog of hapless2 (HAP2) which is a class of membrane fusion protein required for gamete fusion in a range of organisms including *Plasmodium falciparum* (Liu, Tewari et al. 2008). HAP2 labeled protein was exclusively found in male gamonts in *C. parvum* (Tandel, English et al. 2019). In another study, the *C. parvum* transcriptome was elucidated over a 72 h *in vitro* time-course infection with HCT8 cells using semi-quantitative RT-PCR (Mauzy, Enomoto et al. 2012). The 48 h-specific genes from that study were also examined in the 33 RNA-Seq data sets analyzed here (**Supplementary Figure 2**). Both the sex marker genes and 48 h-specific genes show an expression peak at 48 hr. At 48 h, expression levels from batch D samples are much higher than batch C samples. cgd8_2220 is very low/not expressed at 48 h in batch C samples. These results indicate that both batch C and D samples showed the commitment of sexual development, but commitment was more pronounced in batch D. It is possible that sequencing depth could be influencing this difference as the normalization process (VST) between samples tends to reduce the variation of genes with low read support. Interestingly, batch C and D samples used different host cell types (**Table 1**). Batch D *C. parvum* parasites were cultured in HCT-8 (Human intestine cells) while batch C parasites were cultured in MDBK (Bovine kidney cells). Although sexual stages have been observed in MDBK cells (Villacorta, de Graaf et al. 1996), it is possible that the adaptation of *C. parvum* to MDBK cells is lower than HCT-8 cells as hosts. Thus, a slower or lower conversion rate was observed.

### lncRNAs are developmentally regulated

We visualized gene expression profiles across the 33 RNA-Seq samples used in this study (**Table 1**) for both mRNA (**Figure 3A**) and lncRNAs (**Figure 3B**). To identify genes with a similar expression profile, we used the k-mean algorithm to cluster mRNAs and lncRNAs separately. The k value was selected as the smallest value that allowed the separation of genes from different time points while keeping genes from different samples of the same timepoint together despite the batch effects present within each time point. As a result, mRNAs and lncRNAs were clustered into seven and nine broad co-expression groups, respectively.

**Figure 3.**
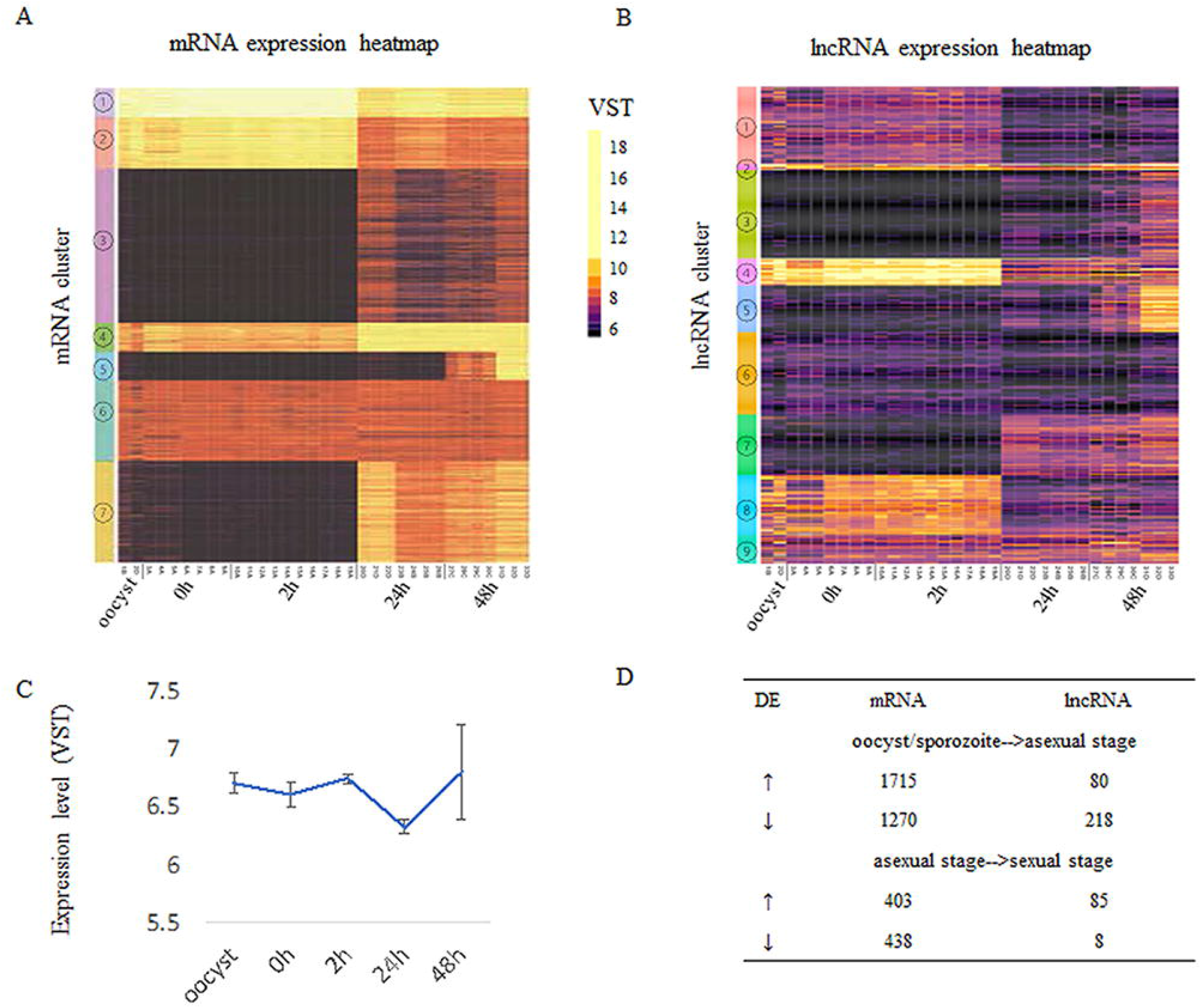
Developmentally regulated lncRNAs. A) Heatmap of mRNA expression across 33 RNA-Seq samples (Table 1). B) Heatmap of lncRNA expression across 33 RNA-Seq samples (Table 1). Expression clusters generated by K-means are indicated by colored bars on the left-most edge. C) The average expression level and standard deviation of lncRNAs at each time point. D) Differentially expressed (DE) genes are compared between develpmental transitions (oocyst/sporozoite stage, 24 h asexual stage and and 48 h sexual stages. The arrows indicate the direction of change in gene expression. Normalized gene expression values are colored as indicated in the scale located between panels A and B with yellow indicating the highest levels.

The expression of mRNAs in the extracellular stages (oocysts and sporozoites from 0h and 2h) showed a similar expression trend that is quite distinct from the latter two stages. For mRNAs, genes from cluster 1 and cluster 2 were more highly expressed in the extracellular stages but still show expression in the intracellular stages when the vast majority of mRNAs are active. This result is consistent with another transcriptome study of *C. parvum*, which used semi-quantitative RT-PCR over a 72 h time course during *in vitro* development (Mauzy, Enomoto et al. 2012). On the contrary, many lncRNAs showed enriched expression in extracellular stages (oocyst, 0 h, 2 h) and some had expression later at the intracellular sexual development stage (48 h) while the asexual stage (24 h) showed the least lncRNA expression. It is noteworthy that both mRNA and lncRNA have gene sets that are specifically turned on at 48 h (mRNA cluster 5 and lncRNA cluster 5). The average expression level of lncRNAs suggests they were more abundant or were actively expressed at the oocyst, 0 h, 2 h and 48 h stages (**Figure 3C**) As was observed in the PCA to assess batch effect (**Figure 1B**) there is increased variation at the 48hr time point. Compared to mRNAs which show more upregulation when transitioning from oocyst/sporozoite to the asexual stage (1715 genes vs 1270 genes), lncRNAs have many more genes downregulated (218 vs 80) (**Figure 3D**). Comparing the asexual stage at 24 h to the sexually activated stage at 48 h, fewer mRNAs showed differential expression, with both having ~400 genes upregulated and downregulated. Very few lncRNAs were downregulated in the transition from the asexual to sexually activated stage but 85 lncRNAs were upregulated. The 85 upregulated lncRNAs did not significantly overlap with the lncRNAs that were downregulated between the extracellular to the asexual stage. Here we only used 48 h samples from batch D to analyze differential expression since this batch has clear sexual stage marker gene expression, as discussed above. The developmentally regulated expression pattern of lncRNAs is indicative of their importance in extracellular and sexual stages. It is important to note that overall, the levels of lncRNA expression are lower than the expression levels observed for mRNAs (Figure 3).

### Correlation of LncRNA expression with neighboring mRNA expression

LncRNA mediated gene regulation can be achieved by various mechanisms (Li, Baptista et al. 2020). One mechanism is transcriptional interference that usually results in repression of the target gene. LncRNAs can also regulate target gene expression through epigenetic mechanisms. Additionally, translational regulation by lncRNA has also been reported. Therefore, to understand the potential roles of lncRNA transcription or transcripts, we studied the correlation between lncRNA and neighboring gene mRNA expression in *C. parvum* (**Supplementary Table 3**). We found that compared to random gene pairs, lncRNAs and their upstream mRNAs have a higher positive correlation of expression level (**Figure 4**) despite the fact that potential read-through transcripts have already been removed. Bidirectional promoters have been reported in many organisms especially species with compact genome sequences. In *Giardia lamblia*, bidirectional transcription is considered to be an inherent feature of promoters and contributes to an abundance of antisense transcripts throughout the genome (Teodorovic, Walls et al. 2007). Thus, some transcriptionally positive correlated lncRNA and upstream mRNAs pairs in *C. parvum* are expected to share bidirectional promoters. However, a large proportion of lncRNA and neighbor mRNAs do not show an apparent expression correlation.

**Figure 4.**
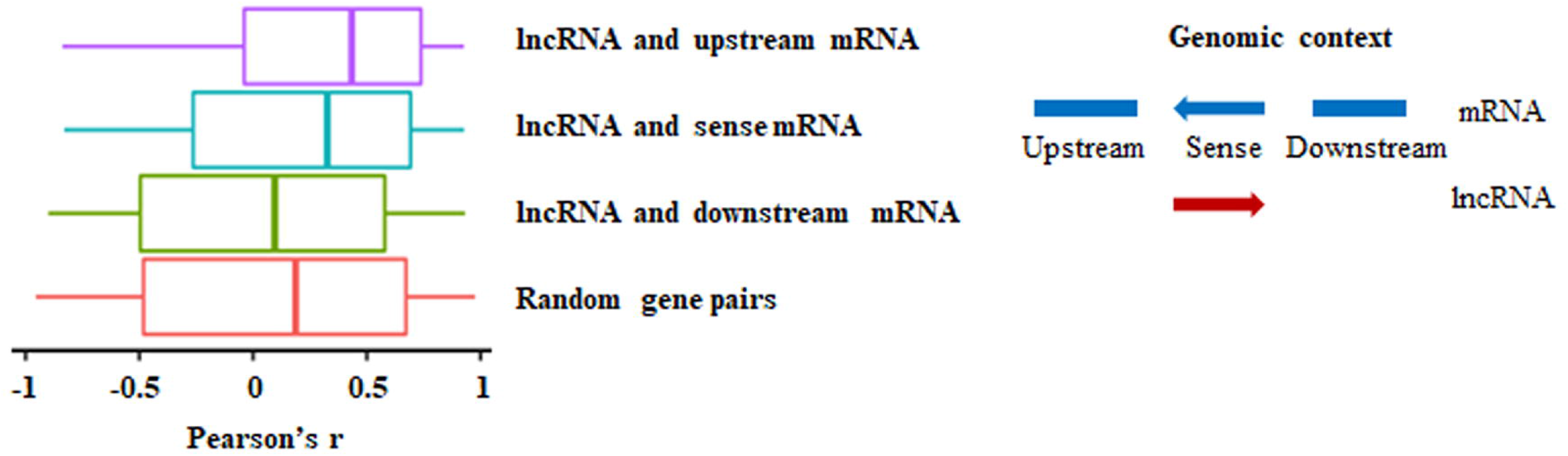
lncRNA expression relative to neighboring mRNAs. The Pearson correlation coefficient was used to measure the expression correlation of different types of gene pair relationships using VST expression levels from the 33 RNA-Seq samples. Random genes pairs were genes that were randomly selected from any lncRNA candidates or mRNA genes. The median value is indicated by a vertical line in each box plot. A graphical representation of the relative position of the mRNA being evaluated to the lncRNA is indicated on the right side. The antisense lncRNA (red arrow) is shown relative to the upstream and downstream mRNAs on the same strand as the sense mRNA.

### Many lncRNAs are conserved between *C. parvum, C. hominis* and *C. baileyi*

Evolutionary conservation of a lncRNA can imply functional importance. lncRNAs can be conserved in different dimensions: the sequence, structure, function, and expression from syntenic loci (Diederichs 2014). lncRNAs are considered to be poorly conserved at the primary sequence level between genera as reported in many higher eukaryotes. Here, we looked for expression conservation of *C. parvum* lncRNA in two other *Cryptosporidium* species from syntenic loci with available stranded RNA-Seq data that include *C. hominis*, a very close relative of *C. parvum* and *C. baileyi*, a distant relative that infects birds (Slapeta 2013). First, we assembled the oocyst/sporozoite transcriptome (the only stranded samples that exist) of *C. hominis* 30976 and *C. baileyi* TAMU-09Q1 by the same methods as used previously except we did not use reference genome annotation guidance (-G). A total of 167 *C. parvum* antisense lncRNAs were detected in both *C. hominis* and *C. baileyi* (**Supplementary Table 4**). Of these, 10 are putatively conserved in *P. falciparum* (**Supplementary Table 5**) based on the presence of antisense lncRNA expression of the orthologus sense gene in *P. falciparum* (Broadbent, Broadbent et al. 2015). No significant sequence similarities were detected among the antisense lncRNAs of these orthologs in *C. parvum* and *P. falciparum*, this is not surprising as little similarity would be found at the level of the sense mRNA’s either given the evolutionary distance and AT bias of *P. falciparum*.

Since the samples of *C. hominis* 30976 and *C. baileyi* TAMU-09Q1 were from oocysts/sporozoites, we focused on 48 of the 167 conserved *C. parvum* lncRNAs that showed a higher expression level in the extracellular stages (**Figure 5**). The corresponding sense mRNAs were involved in various biological processes. Translation related functions, including translational initiation (cgd7_2430, translational initiation factor eIF-5) and protein folding (cgd2_1800, DnaJ domain-containing protein), were seen in the sense mRNAs. A positive correlation of 0.78 and a negative correlation of −0.78 was calculated for cgd7_2430 sense-antisense pair and cgd2_1800 sense-antisense pair, respectively. In addition, a few mRNAs that encode putative secreted proteins (cgd5_10 and cgd4_3550) also showed a high positive correlation of expression with the corresponding antisense.

**Figure 5.**
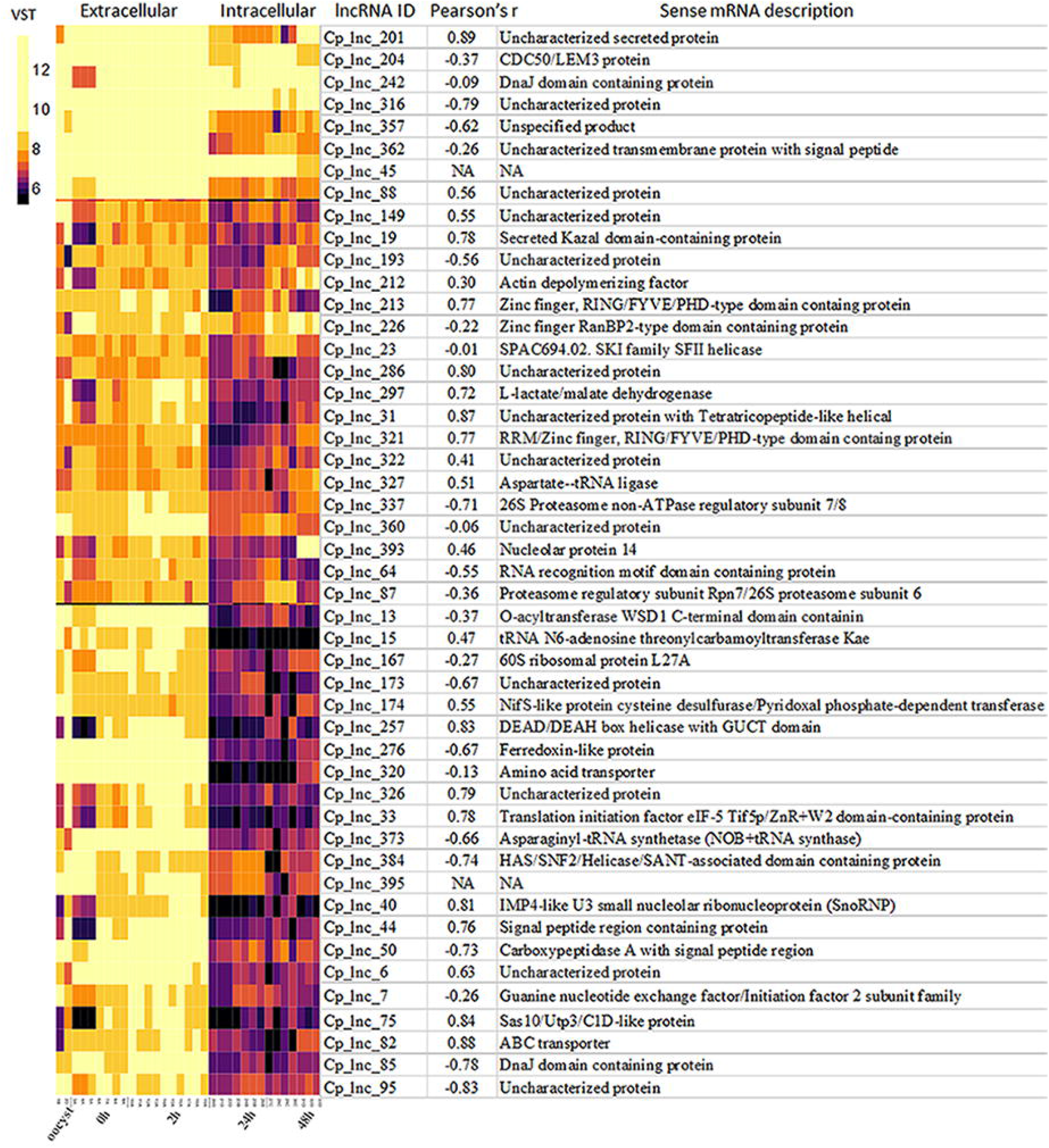
Highly expressed *C. parvum* lncRNAs with conservation and expression in *C. hominis* and *C. baileyi* oocysts. The heatmap visualizes the lncRNA expression level of 48 conserved *C. parvum* genes across 33 RNA-Seq samples, grouped as extracellular (oocyst/sporozoite, 0 h and 2 h) and intracellular (24 h and 48 h) stages. The lncRNA name and the corresponding sense mRNA description are listed. An mRNA description of “NA” indicates the lncRNA is intergenic. The sense-antisense expression correlation coefficient is shown in the bracket. The color scale is shown on the left. Yellow indicates high levels of expression.

### lncRNA prediction validation

In the RNA-Seq data, Cp_lnc_51 was expressed in oocyst/sporozoites while the associated sense mRNA cgd1_380 (Ubiquinone biosynthesis protein COQ4) was seen to be silenced. The expression levels for each were validated by stranded RT-qPCR (**Figure 6A**). To validate lncRNA by strand-specific RT-PCR (Ho, Donaldson et al. 2010), a specific RT primer was designed for each gene to generate the cDNA with strand information retained (**Supplementary Table 2**). Cp_lnc_51 contains an intron. The splicing of Cp_lnc_51 was confirmed by RT-PCR and agarose gel electrophoresis to assess the transcript size (**Figure 6B**). We randomly selected additional five lncRNAs for validation. Two out of five, Cp_lnc_82 (**Figure 6C**) and Cp_lnc_93 (**Figure 6D**) were validated by qPCR. The relative expression of sense-antisense is consistent with the RNA-Seq data.

**Figure 6.**
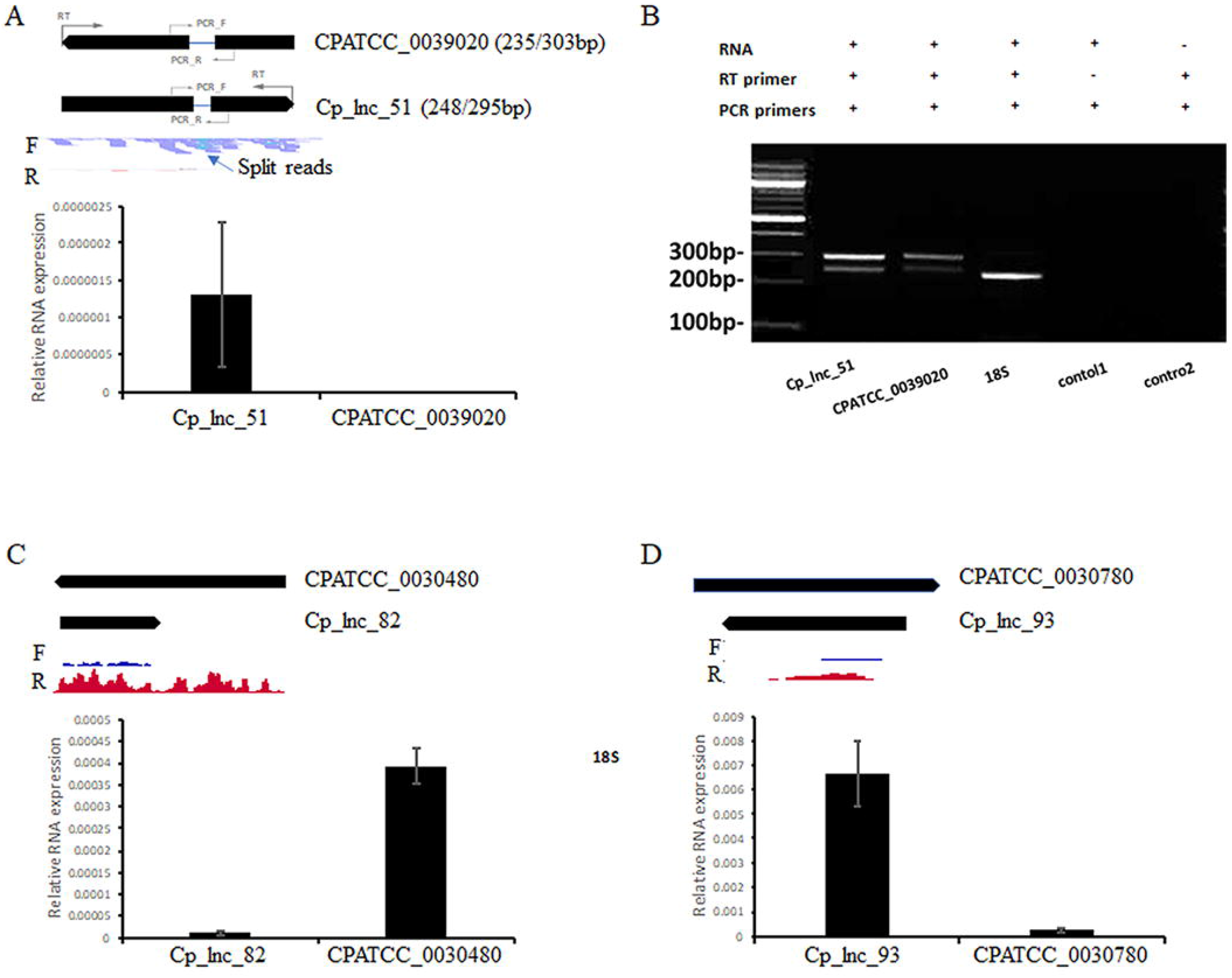
lncRNA candidates expression and intron structure validation. A) The expression level of lncRNA Cp_lnc_51 and the corresponding sense mRNA CPATCC_0039020 validated by RT-qPCR, the expression was normalized to 18S. The annotated genome model is shown on the top with RNA-Seq reads mapped to the genomic region. Location of RT primers and PCR primers for each gene are shown with the gene models. Reads are separated by the mapped strand: forward strand (F) and reverse strand (R). The intron structure of Cp_lnc_51 is indicated by the split reads. B) Splicing of the Cp_lnc_51 transcript is supported by the various length of transcripts from intron splicing, shown on agarose gel with the expected size. The expected size of the products with/without intron is indicated next to the gene name in panel A. 18S is used as positive control with expected size of 239bp. Control 1 is negative control without RT primer but only PCR primers of CPATCC_0039020 added. Control 2 is also negative control with both RT and PCR primers of CPATCC_0039020 but no RNA template added. C) The expression level of lncRNA Cp_lnc_82 and the corresponding sense mRNA CPATCC_0030480 validated by RT-qPCR, the expression was normalized to 18S. D) The expression level of lncRNA Cp_lnc_93 and the corresponding sense mRNA CPATCC_0030780 validated by RT-qPCR, the expression was normalized to 18S. The RNA-Seq coverage in C and D is with range 0-100 CPM (counts per million reads mapped).

## Discussion

In this study, we utilized stranded RNA-Seq data from multiple time points during parasite development to systematically identify and characterize lncRNAs in *C. parvum*. 396 high-confidence lncRNAs were identified, 363 occur as antisense transcripts to mRNAs and 33 are encoded in intergenic locations. Nearly 10% of predicted mRNAs are covered by an antisense lncRNA. This pervasive antisense transcription suggests an important function of lncRNA in *C. parvum*. The lncRNAs were analyzed to determine expression profiles, promoter motifs for coordinately expressed transcripts, transcriptional relationships with upstream and downstream mRNAs and conservation among three *Cryptosporidium* species with stranded RNA-Seq data available.

To investigate the expression relationship of lncRNAs and their neighboring mRNA encoding genes, we calculated the expression correlation of different type of gene pairs by Pearson coefficient and noticed a higher positive correlation of expression between lncRNA and its upstream mRNAs compared to random gene pairs. Many sense and antisense pairs also showed a positive correlation of expression. Notably, spurious correlations of gene expression can happen if the biological variation among samples is too large. Due to the challenge of *in vitro* culture for *C. parvum* and very low volume, hence number of the parasite transcripts compared to their host cells, samples from the early intracellular stages are rare and usually contain very low levels of parasite transcripts. In this study, the transcriptome data from early intracellular stages was absent. We detected a bimodal shape for the distribution of expression correlation with random gene pairs showing trends at both high positive and negative values, probably due to spurious correlation. However, lncRNAs showed much less negative correlation of expression with both upstream and corresponding sense mRNA than random gene pairs. Thus, the higher positive correlation between lncRNA and the neighboring mRNAs may suggest pervasive bidirectional promoters in *C. parvum*. Another possibility is that lncRNAs function as positive regulators of the neighboring mRNA expression. In *P. falciparum*, ncRNAs derived from GC-rich elements that are interspersed among the internal chromosomal *var* gene clusters are hypothesized to play a role in *var* gene activation while the mechanism is unclear (Guizetti, Barcons-Simon et al. 2016, Barcons-Simon, Cordon-Obras et al. 2020). lncRNAs have been associated with chromatin remodeling to achieve transcriptional regulation in many studies (Li, Baptista et al. 2020). One example is that a lncRNA HOTTIP transcribed from the 5’ tip of the HOXA locus that coordinates the activation of HOXA genes by maintaining active chromatin (Wang, Yang et al. 2011).

Functional enrichment analysis is challenging for this parasite due to the large number of uncharacterized proteins and incomplete pathways (Rider and Zhu 2010). Functional analysis for antisense associated sense mRNA by GO enrichment didn’t significantly define key biological processes. However, the enriched expression of lncRNA in extracellular stages and late intracellular stages, the time point when the parasite starts to have sexual commitment and produce gametes, suggests potential critical roles that lncRNAs may play during these life cycle stages. One possibility is that lncRNAs are involved in transcriptome pre-loading in macrogamont that will eventually become an oocyst. It is also possible that lncRNA play a role in transcriptional regulation or that antisense lncRNAs may play roles in the post-transcriptional process. It has been reported that lncRNAs can regulate translation by stabilizing mRNAs, triggering mRNA degradation or triggering translation process by interactions with associated machineries (Li, Baptista et al. 2020). As shown in this study, antisense transcripts have a strong bias towards covering the 3’ end of the sense mRNA. This property has also been reported in other organisms, including the malaria pathogen *P. falciparum* (Siegel, Hon et al. 2014, Broadbent, Broadbent et al. 2015). As these authors suggest, one possibility is that antisense RNAs arise from promiscuous transcription initiation from nucleosome depleted regions (Siegel, Hon et al. 2014). It is also known that the 3’ UTR of mRNAs can contain elements that are important for transcript cleavage, stability, translation and mRNA localization. The 3’ UTR serves as a binding site for numerous regulatory elements including proteins and microRNAs (Jia, Yao et al. 2013, Tushev, Glock et al. 2018). In humans, the antisense *KAT5* gene has been reported to promoted the usage of distal polyA (pA) site in the sense gene *RNASEH2C*, which generated a longer 3’ untranslated region (3’ UTR) and produced less protein, accompanied by slowed cell growth (Shen, Li et al. 2018). Whether the 3’ end bias of antisense expression related to its function and translation repressor need further investigation. One future direction is to take advantage of single-cell sequencing approaches and look at the transcriptomic details of male and female gametocytes. It will be interesting to see if lncRNAs are specifically expressed in male and female gametocytes and whether or not some lncRNAs are restricted specifically to these gamonts or if they are also detected elsewhere, e.g., in oocysts where they could, perhaps, have a role in transcriptional or post-transcriptional gene regulation, or mRNA stability. The amount of active transcription as opposed to RNA pre-loading in the female gamont (future oocyst) is not known.

Twenty-two *C. parvum* lncRNAs have been detected in the host cell nucleus (Wang, Gong et al. 2017). Of these, 18 were detected in this study. Motif analysis was conducted on the exported lncRNA transcript sequences but no significant similarity or motif was detected relative to the larger pool of lncRNA candidates identified in this study. This raises the question of what signal is responsible for lncRNA export. Further studies are needed.

A significant roadblock in lncRNA research is the determination of their function. Genetics studies are particularly tricky with anti-sense transcripts of the sequences overlap coding sequences closely, as they do in *C. parvum* because genetic alterations of the sequence affect the sense and anti-sense transcripts. lncRNAs with similar functions often lack sequence similarity (Kirk, Kim et al. 2018). Many known lncRNAs function by interacting with proteins. Proteins often bind RNA through short motifs (three to eight bp in length) (Ray, Kazan et al. 2013). It was hypothesized that lncRNAs with shared functions should harbor motif composition similarities (Kirk, Kim et al. 2018). In this study, nucleotide composition of lncRNAs varies between those that are antisense and those that are encoded in intergenic regions. We see many lncRNAs have higher CT-rich content compared to mRNAs (**Supplementary Figure 3**). Interestingly, lncRNAs can be grouped into CT-rich and AT-rich, with most of the intergenic lncRNAs belonging to the AT-rich group. The difference of nucleotide composition gives rise to the speculation that the machineries interacting with antisense and intergenic lncRNAs may be different in *C. parvum*.

lncRNA prediction using short reads in organisms with compact genome sequences, including *C. parvum* is limited due to the difficulty of separating independent lncRNA transcription from neighboring transcriptional read-through noise. In this study, we used a customized pipeline with strict criteria designed to minimize false positives from background noise such as transcriptional read-through. Many antisense transcriptions cover all or most of the sense mRNA transcript. To further improve the discovery of full-length lncRNA and any isoforms, long-read approaches such as Iso-Seq (Pacific Biosystems) and single molecule pore-sequencing approaches [Oxford Nanopore Technologies (ONT)] are needed. Although obtaining sufficient high-quality RNA from intracellular stages is still challenging, hybrid capture approaches (Gnirke, Melnikov et al. 2009, Amorim-Vaz, Tran Vdu et al. 2015) can be utilized to obtain *Crptosporidium* RNA to be used for, direct RNA sequencing on the ONT platform providing additional insights into the RNA biology of *Crypptosporidium*. Besides, long-read sequencing would also enable better annotation of mRNA UTR boundaries (Chappell, Ross et al. 2020), which can be used to further investigate the 3’ UTR bias of antisense transcription as observed in this study.

It is important to understand how species evolve and adapt to their environment. Due to the poor conservation of lncRNA reported in higher eukaryotes (Johnsson, Lipovich et al. 2014) and the large phylogenetic distance among *Cryptosporidium* species (Slapeta 2013), it is noteworthy that many lncRNAs detected in *C. parvum* were also seen expressed in *C. baileyi*. It indicates that RNA regulation could be a common and critical strategy for *Cryptosporidium* gene regulation or interactions with their hosts. The discovery of conserved antisense lncRNA expression between *C. parvum* and *P. falciparum* orthologs revealed that many important mRNAs have antisense expression including a methyltransferase protein, a palmitoyltransferase and a copper transporter. lncRNAs function by interaction with DNA, RNA or proteins. Thus, the structure of lncRNAs could be under stronger selection than their sequence. Since most lncRNAs are antisense in *C. parvum*, to separate conservation of lncRNA from the conservation of mRNA sequence could provide further insights into lncRNA evolution. Selection pressures that independently act to maintain sequence and secondary structure features can lead to incongruent conservation of sequence and structure. As a consequence, it is possible that analogous base pairs no longer correspond to homologous sequence positions. Thus, possible selection pressures independently acting on sequence and structure should be taken into account (Nowick, Walter Costa et al. 2019). Despite the increasing acknowledgment that ncRNAs are functional, tests for ncRNAs under either positive or negative selective did not exist until recently (Walter Costa, Honer Zu Siederdissen et al. 2019). This type of analysis will assist in identifying candidates to prioritize for further functional lncRNA investigations.

## Supporting information

lncRNA fasta sequence file

