## Supplementary Figures for "Stage-Specific Long Non-coding RNAs in *Cryptosporidium parvum* as Revealed by Stranded RNA-Seq"

Supplementary Material

The figure legends are required to have the same font as the main text, 12 point normal Times New Roman, single spaced. Please use a single paragraph for each legend and prepare the figures keeping in mind the PDF layout.

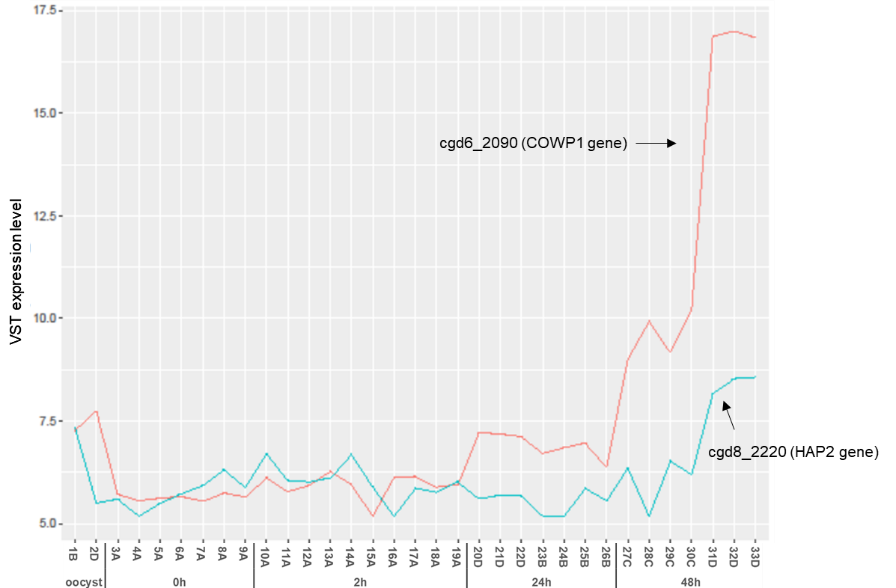

**Supplementary Figure 1.** Expression profile of sex-specific marker genes. VST normalized expression levels of female marker COWP1 gene and male marker HAP2 gene in the 33 RNA-Seq samples are visualized. The sample ID is as indicated on the x-axis.

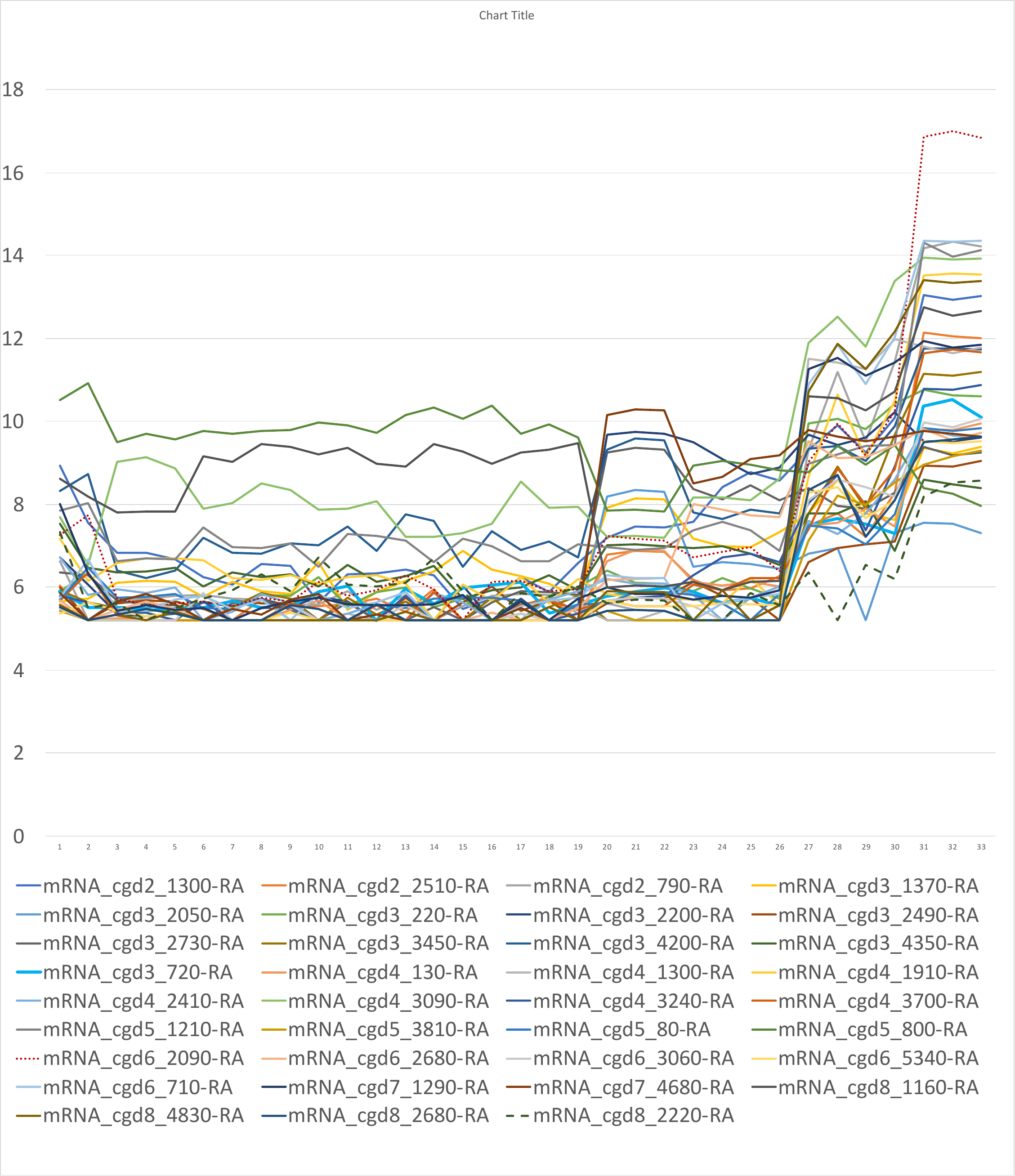

**Supplementary Figure 2.**  **Expression profile of 48 h-specific genes.** VST normalized expression levels of 48 h-specific genes selected from the literature are visualized within the 33 RNA-Seq samples. The sample ID is as indicated on the x-axis.

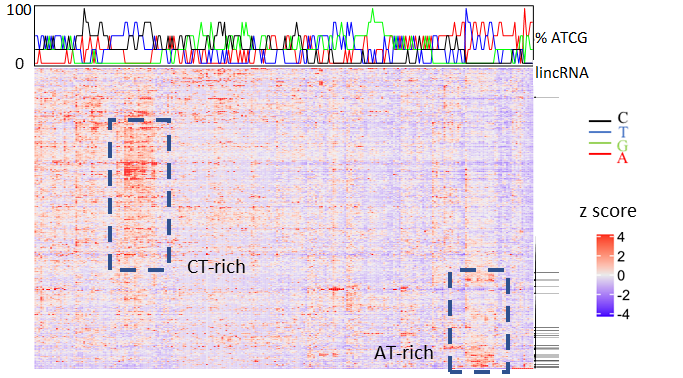

**Supplementary Figure 3**. **Relative 4-mer abundance profile of lncRNAs.** Each row represents a lncRNA candidate, and each column represents a 4-mer (4^4^=256 4-mers). The abundance of each 4-mer is indicated by color. Z score is calculated based on relative abundance with respect to mRNAs. Red suggests the 4-mer in lncRNAs is more adunacnt than in mRNAs. Blue indicates the opposite. The intergenic lncRNAs (lincRNAs) are indicated by the black line on the right , the remainder are antisense transcripts. The top panel shows the percent of ATCG for each 4-mer. From k-mer size 4 to 8, the clustering pattern remained but was less apparent. Thus, only results from 4-mer are shown here.
